# An in vitro and in vivo approach for the isolation of Omicron variant from human clinical specimens

**DOI:** 10.1101/2022.01.02.474750

**Authors:** Pragya D. Yadav, Nivedita Gupta, Varsha Potdar, Sreelekshmy Mohandas, Rima R. Sahay, Prasad Sarkale, Anita M. Shete, Alpana Razdan, Deepak Y. Patil, Dimpal A. Nyayanit, Yash Joshi, Savita Patil, Triparna Majumdar, Hitesh Dighe, Bharti Malhotra, Jayanthi Shastri, Priya Abraham

## Abstract

Due to failure of virus isolation of Omicron variant in Vero CCL-81 from the clinical specimen’s of COVID-19 cases, we infected Syrian hamsters and then passage into Vero CCL-81 cells. The Omicron sequences were studied to assess if hamster could incorporate any mutation to changes its susceptibility. L212C mutation, Tyrosine 69 deletion, and C25000T nucleotide change in spike gene and absence of V17I mutation in E gene was observed in sequences of hamster passage unlike human clinical specimen and Vero CCL-81 passages. No change was observed in the furin cleavage site in any of the specimen sequence which suggests usefulness of these isolates in future studies.

## Introduction

The ability of SARS-CoV-2 to rapidly mutate has been the biggest challenge the world has faced while responding to the pandemic. Speculation and the anticipation of SARS-CoV-2 becoming endemic has caome to an end with the recent emergence of a heavily mutated variant, Omicron (B.1.1.529). The fifth variant of concern (VOC) was first reported by South Africa followed by Botswana and Hong Kong in November 2021.^1^ The variant has now spread to all six continents in a few weeks’ time^2^ with India reporting more than 600 cases to date. Omicron has posed a serious public health concern due to the mutations/deletions associated with increased binding affinity to ACE2 (S:Q498R, S:N501Y), increased transmissibility (S:H655Y, S:N679K, S:P681H), increased viral load (N:R203K, N:G204R), innate immune evasion (ORF1a:L3674-, ORF1a:S3675-, ORF1a:G3676), and S-gene target failure (S:H69-).^3^

The global scientific community is putting the best efforts to determine the susceptibility, transmissibility, immune escape and severity associated with Omicron. Such an emergency demands the availability of Omicron virus isolate. Many variants of SARS-CoV-2 have been isolated successfully utilizing Vero, Huh7, and human airway epithelial cells. Various cell lines such as Vero CCL-81, Vero SLAM, MA104, BGM and Caco-2 have also supported the replication of SARS-CoV-2.^4^ However, there are very limited studies that have reported the isolation of SARS-CoV-2. Here, we report the isolation and characterization of the Omicron variant from clinical specimens of two COVID-19 cases using the Syrian hamster model and Vero CCL-81 cells.

## Materials and methods

The oropharyngeal/nasopharyngeal swabs of SARS-CoV-2 positive (E gene: Ct value range 17-30) international travelers (n=73) and contact case (n=1) from Delhi and Mumbai, India were collected during 10th October to 13th December 2021. These specimens were transported under cold chain and inoculated onto Vero CCL-81 which was maintained in Eagle’s minimum essential medium (MEM; Gibco, UK) supplemented with 10 per cent foetal bovine serum (FBS) (HiMedia, Mumbai), penicillin (100 U/ml) and streptomycin (100 mg/ml). Likewise, 100 μl was inoculated onto 24-well cell culture monolayers of Vero CCL-81, before growth medium was decanted. The cells were incubated for one hour at 37°C to allow virus adsorption, with rocking every 10 min for uniform virus distribution. After the incubation, the inoculums specimen was removed and the cells were washed with 1X phosphate-buffered saline (PBS). The MEM supplemented with two per cent FBS was added to each well. The cultures were incubated further in five per cent CO2 incubator at 37°C and observed daily for CPEs under an inverted microscope (Nikon, Eclipse Ti, Japan). Cellular morphological changes were recorded using a camera (Nikon, Japan). Cultures that showed CPE on PID-5 were centrifuged at 4815 × g for 10 min at 4°C; the supernatants were processed immediately or stored at −86°C. Further, those that showed CPE were grown in T-25 flasks at P-2 and titration was done after serial dilution. Tissue culture infective dose 50 per cent (TCID50) values were calculated by the Reed and Muench method. The cells were examined microscopically for cellular morphological changes following inoculation.^5^

Simultaneously, the Next-generation sequencing for clinical specimens, hamster specimens and virus isolate was performed on the Illumina MiniSeq Machine. Briefly, the ribosomal RNA depletion was performed using Nebnext rRNA depletion kit (Human/mouse/rat) followed by cDNA synthesis using the first strand and second synthesis kit. The RNA libraries were prepared using TruSeq Stranded Total RNA library preparation kit. The amplified RNA libraries were quantified and loaded on the Illumina sequencing platform after normalization. Reference-based mapping was performed to retrieve the complete genome sequence of the clinical specimens, hamster specimens and using the CLC genome workbench V 20.0.4 and submitted to the public repository i.e., GISAID. A phylogenetic tree was generated using the MEGA software version 7. The nucleotide variation of the sequences analyzed in the study was generated using the highlighter plot (https://www.hiv.lanl.gov/cgi-bin/HIGHLIGHT/highlighter.cgi).

With subsequent efforts, two Omicron positive specimens with [MCL-21-H-11828 (group-A) Ct: 20 and MCL-21-H-12521/1 (group-B); Ct: 25] identified by whole genome sequencing were selected for in vivo virus isolation. The Syrian hamsters (8-10 weeks old) were anesthetized using the Isoflurane. A hundred microlitre volumes of each specimen were inoculated through the intranasal route into two hamsters. The nasal turbinate (NT), lung specimens of passage 1 (P1) were further paasaged into new batch of hamsters (P2). The hamsters were sacrificed and NT, lung specimens were collected on the 3^rd^ day post-infection (DPI). NT scrapings and lung tissues (20% suspension) were prepared in sterile Minimum Essential Medium (MEM; Gibco, USA) using a homogenizer and were screened using rRT-PCR.^7^ SARS-CoV-2 positive NT and lung homogenized suspension of hamsters (P1 and P2) were further inoculated onto Vero CCL-81 cells.

## Results

An attempt of isolation using Vero CCL-81 yielded 10 Delta derivatives [AY.125 (n=4), AY.122 (n=2), AY.89 (n=1), AY.46 (n=1), AY.4 (n=1), AY.4.2.1 (n=1)]. The Omicron positive clinical specimens were blindly passaged thrice; however, replicating virus couldn’t be detected using rRT-PCR and no CPE was observed.^5,7^

The complete genome (>98%) were retrieved from 61 clinical specimens. Forty six sequences belonged to Delta derivatives [Delta (n=2), B.1.617.2.106 (n=1), B.1.617.2.120 (n=1), B.1.617.2.122 (n=2), B.1.617.2.125 (n=3), B.1.617.2.4 (n=2), B.1.617.2.43 (n=1), B.1.617.2.6 (n=2), B.1.617.2.85 (n=2), AY.122 (n=11), AY.125 (n=4), AY.36 (n=1), AY.4 (n=4), AY.4.2.1 (n=1), AY.43 (n=4), AY.46 (n=1), AY.5.4 (n=1), AY.89 (n=1), AY.9.2 (n=2)]; while 15 were identified as B.1.1529. Among the 15 Omicron cases (median age:34 years, male:6/female:9), 14 were international travelers from United Arab Emirates, South Africa, Nigeria, Tanzania, France, United Kingdom, Germany, Netherland and one contact case from Mumbai. Thirteen were asymptomatic while two had mild cold, cough and body ache.

In vivo isolation of the virus from clinical specimens demonstrated active replication of the virus was observed in both the hamsters of group-A, while group-B hamsters showed replication in NT but not in lungs (Figure 1A). NT and lungs of hamsters group-A had genomic viral RNA (gRNA) copies of 2.0–2.2×10^9^ and 2.3×10^5^–1.9×10^8^ per ml respectively (Figure 1 B). While, NT of hamsters from the group-B had gRNA copies of 1.5×10^8^–8.3×10^9^. The P1 NT and lung specimens of group-A hamsters inoculated in new batch of hamsters (P2) demonstrated gRNA copies of 2.6×10^10^ and 6.6×10^10^ in NT and lungs respectively on 3^rd^ DPI (Figure 1B).

**Figure 1:**
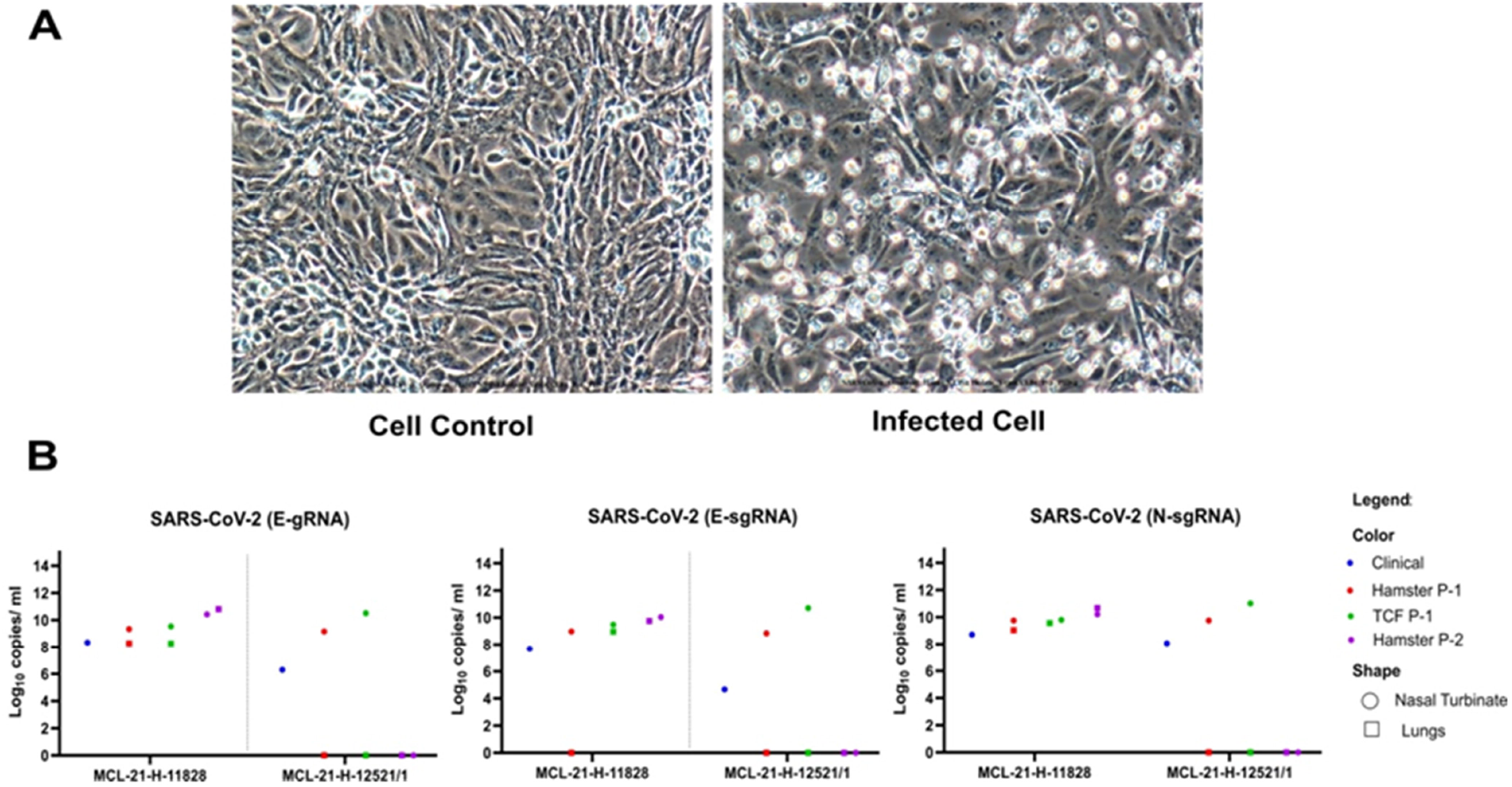
Omicron variant virus isolation in the Vero CCL-81 Cells and their viral titer: A) Vero CCL-81 cell control (Left) and Omicron infected Vero CCL-81 cell (Right) at passage 2, PID-4. B) The viral load of the SARS-CoV-2 in human clinical samples, hamster specimens at passage 1 and 2 and Vero CCL-81 at passage 1 culture. The genomic RNA and subgenomic RNA load for the E gene along with N gene from nasal turbinate and lungs of two different clinical specimens.

The hamsters NT and lung homogenized specimens (P-1) inoculated onto Vero CCL-81 cells (P-1) showed evidence of cell rounding and detachment from post infection day (PID)-4 in P-1. Syncytial cells formed large cell masses that increased in size and number as the infection progressed. Enhanced CPE was noted in P-2 at PID-3. Incomplete cell detachment from the tissue culture plate surfaces, rounding and refractive appearance of infected cells was observed (Figure 1A). The hamster NT and lungs specimens of group-A showed typical CPE in P-1 and P-2 of Vero CCL-81 with gRNA copies of 3.4×10^9^ to 1.9×10^10^ and 1.8×10^8^ to 1.0×10^9^ per ml respectively. Sub genomic RNA was also detected in isolates obtained from NT specimens of two hamsters (P1: 2.9×10^9^ and 5.0×10^10^; P2: 4.5×10^9^ and 1.3×10^10^) and lung specimen of one of the hamster (P1: 1.8×10^8^; P2: 1.1×10^9^) [Figure 1B]. While only hamster NT (P1 and P2) specimen of group-B displayed CPE in Vero CCL-81 with gRNA copies of 3.3×10^10^ and 4.8×10^10^ (Figure 1 B).

The molecular characterization of the virus isolates obtained with in vivo and in vitro method confirmed them as Omicron (Figure 2A). The Omicron variants and its isolates clustered together with other representative SARS-CoV-2 sequences (Figure 2B). Single nucleotide variations (SNV) were observed using the CLC genomics Workbench ver. 22.0. The differences were observed in the Omicron sequences from the human clinical, hamster and Vero CCL-81 samples. The insertion of AGT nucleotides at genomic position 22195 was observed in the hamster P1 sequence and this insertion changed two amino acid encoding codons; (AG) lied in the first codon while the third nucleotide insertion ‘T’ lied in the second codon. Hence the reading frames of the codons were changed from ATA to AAGTGC. The tyrosine at 69 amino acid position of spike was lost in the hamster in P1 and observed in hamster P2. These mutations further needs to be explored which might have facilitated the viral entry and replication in hamster. Further a nucleotide change of ‘C’ to ‘T’ was observed at position 25000 only in the hamster sequences.

**Figure 2:**
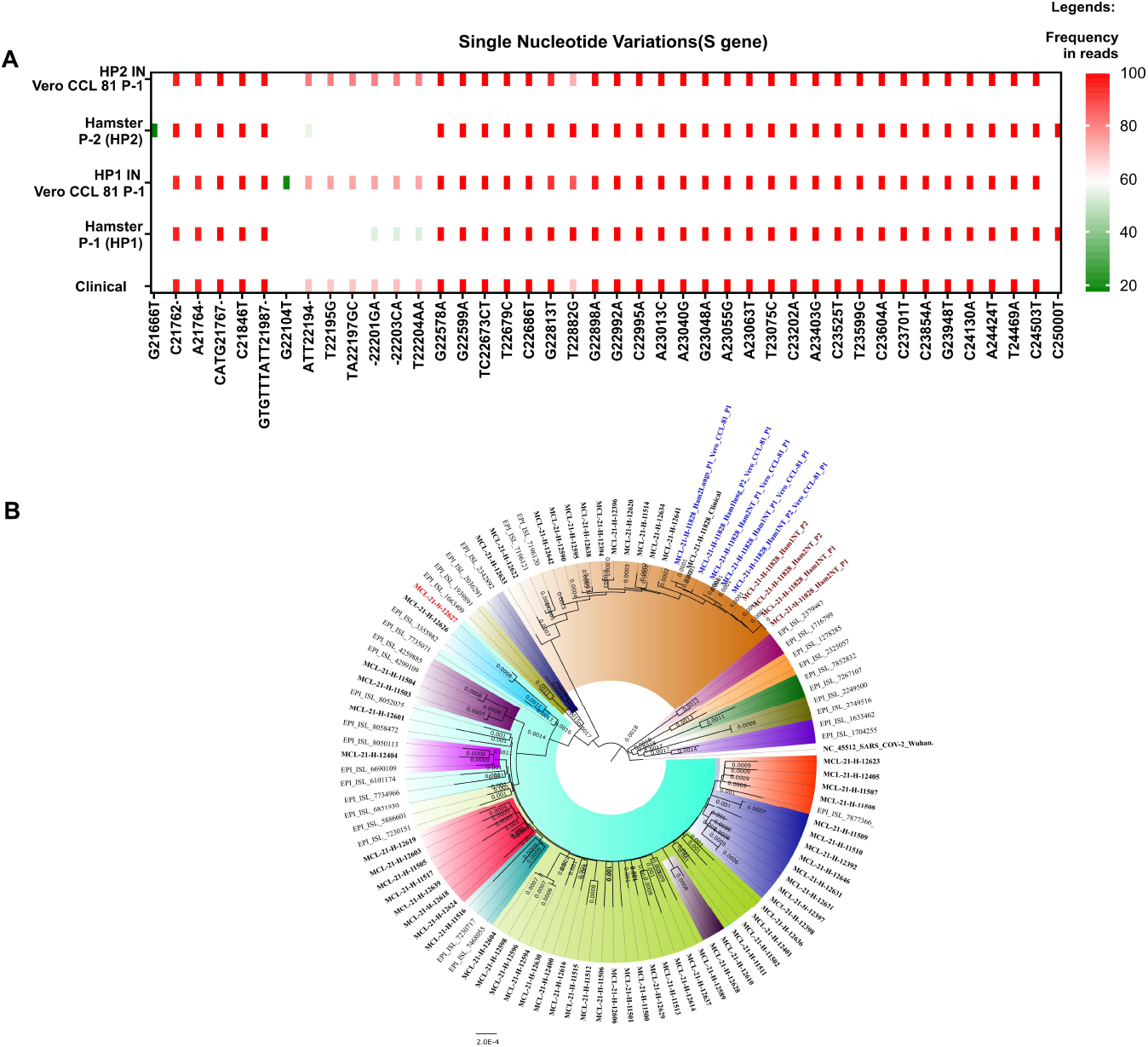
SNV and phylogenetic tree of the Omicron clinical and its isolate sequences: **A)** Single nucleotide variation in the S gene and its frequency in different in vivo and in vitro sequences. The x axis is the SNVs and Y axis marks the frequency. B) Neighbour joining tree with a bootstrap replication of 1000 cycles to assess statistical robustness. Different SARS-CoV-2 variants are marked in various colors.

A gain of an SNV expand was observed at the E gene [G26293A (V17I)] of P-1 Vero CCL-81 isolate, while it was absent in clinical as well as hamster sequences. Further an SNV was also observed at M gene [C26577G (Q19E)] of P1 hamster which was not observed in clinical sample, Vero CCL-81 P1 and hamster P2. An in-depth analysis would be required to understand the gain or loss of this specific mutation.

The frequency of the N440K amino acid change was observed in the hamster sequences (100%) and in the human clinical sequences (68.5%). The Vero CCL-81 passage demonstrated the decrease by 86.0% in Vero CCL-81 P1 of the hamster (P1) and 71.79% in Vero CCL-81 (P1) of the hamster (P2). The variation in the frequency of the N440K in human clinical specimens, hamster specimens and Vero CCL-81 passages could be due to the adaptive mutations in different host species. A recent study linked this mutation to be associated with higher infectious titer.^8^ Further the studies have reported that the N440K strains have capability of escaping the neutralization.^9^

## Discussion

This study reports the isolation of the Omicron variant using the in vivo followed by in vitro method of virus culture. Until now, there is no literature available on the isolation of SARS-CoV-2 using initial in vivo and subsequent in vitro approach. Spike region of the hamster revealed L211C amino acid change along with the C25000T nucleotide change compared to clinical specimen. A percent nucleotide identity of 0.0066% was observed, indicating a minimal change of Omicron sequence in Vero CCL-81 P-1 with hamster P2. However a detailed analysis for further passages with Vero CCL-81 needs to be explored.

Apparently, the sequences of human clinical specimens, hamster specimens and Vero CCL-81 passages didn’t show any mutations in the furin cleavage site during sequential passages in different host species. Such virus isolate would retain its pathogenicity as per the original specimen/isolate. The isolated Omicron virus would be useful in new vaccine development, vaccine efficacy studies, pathogenicity in hamsters, antiviral research and estimation of virus neutralizing titers in vaccinated and recovered individuals.

## Ethical approval

The study was approved by the Institutional Biosafety Committee and Institutional Human Ethics Committee of ICMR-NIV, Pune, India under the project titled ‘Molecular epidemiological analysis of SARS-COV-2 circulating in different regions of India’.

## Author contributions

PDY contributed to study design, data analysis, interpretation and writing. VP, AMS, RRS, SM, AR, DYP, DAN, PS, SP, TM, HD and YJ contributed to data collection, data analysis, interpretation and writing. PDY, DYP, DAN, RRS, AMS, SM, BM, JS, NG and PA contributed to the critical review and finalization of the manuscript.

## Acknowledgement

The author gratefully acknowledges the contribution of Ms. Manisha Dudhmal, Ms. Pranita Gawande, Mrs. Kaumudi Kalele and Ms. Jyoti Yemul from Maximum Containment Facility, ICMR-NIV, Pune.

